# Virulence potential of a multidrug-resistant *Escherichia coli* strain belonging to the emerging clonal group ST101-B1 isolated from bloodstream infection

**DOI:** 10.1101/797928

**Authors:** Ana Carolina de Mello Santos, Rosa Maria Silva, Tiago Barcelos Valiatti, Fernanda Fernandes dos Santos, José Francisco Santos-Neto, Rodrigo Cayô, Ana Paula Streling, Carolina Silva Nodari, Ana Cristina Gales, Milton Yutaka Nishiyama-Junior, Eneas Carvalho, Tânia Aparecida Tardelli Gomes

**Author notes:** corresponding author (ACMS).

## Abstract

*Escherichia coli* EC121 is a multidrug-resistant (MDR) strain isolated from bloodstream infection of an inpatient with persistent gastroenteritis and Zone T lymphoma, that died due to septic shock. Despite causing an extraintestinal infection, it harbors few known virulence factors and was assigned into phylogenetic group B1. To evaluate if the EC121 was pathogenic or opportunistic, its genome was sequenced, and an *in vitro* characterization of some pathogenicity-associated properties was performed. The data retrieved from genome analysis showed that *E. coli* strain EC121 belongs to the O154:H25 serotype, and to ST101-B1, which was epidemiologically linked to extraintestinal infections and antimicrobial resistance spread as well. Moreover, it was closely related to Shiga-toxin producing *E. coli* (STEC). Besides, strain EC121 is an MDR strain harboring 14 antimicrobial resistance genes, including *bla*_CTX-M-2_, and more than 50 complete virulence genetic clusters, which are reported to be associated either with DEC or ExPEC. The strain also displays the capacity to adhere to a variety of cell lineages, and invade T24 bladder cells, as well as the ability to form biofilms on abiotic surfaces, and survive the bactericidal serum complement activity. Additionally, it is virulent in the *Galleria mellonella* model. Altogether, *E. coli* EC121 unveiled to be a pathogen powered by its multi-drug resistance characteristic. Carry out studies providing accurate information about the virulence potential of all kinds of MDR strains are essential because these studies will help in the development of alternative therapies of infection management and spread control of MDR strains.

**Authors summary:** The phylogenetic origin of extraintestinal pathogenic Escherichia coli is mostly associated with phylogroup B2, and the majority of the studies regarding extraintestinal infection focus on the most virulent strains, which might also present multidrug-resistant (MDR) phenotype. Strains belonging to phylogroup B1 and isolated from extraintestinal infections are considered as opportunist pathogens and have their virulence neglected. We focus our study in one MDR strain isolated from bloodstream infection that belongs to phylogenetic group B1 to enlarge the knowledge about the virulence of this kind of strain. We demonstrated that the EC121 is capable of adheres to intestinal and bladder human cells, and invades the latter one; it survives to human serum bactericidal effects and produces biofilm. Additionally, the in vivo assay confirmed the EC121 virulence, showing that it should be considered a pathogenic strain. The genetic analyzes highlighted important aspects of EC121 which are typical from strains of sequence type 101, like its involvement in the spread of antimicrobial resistance genes and its relationship with extraintestinal infection from diverse sources. Information concerning the virulence of MDR strains is important for the development of global actions treating the spread of antimicrobial resistance, as well as to elucidate the pathogenesis of strains that were considered as an opportunist.

## Introduction

*Escherichia coli* is one of the most frequent pathogens isolated from bloodstream infections (BSI) around the world [1–5]. Despite all knowledge of extraintestinal infections due to pathogenic *E. coli,* the number of severe infections and outbreaks caused by these pathogens is rising [2, 6]. Moreover, many of these infections are caused by multidrug-resistant (MDR) strains, leading to a higher burden of disease [7–10]. The term extraintestinal pathogenic *Escherichia coli* (ExPEC) is used to define strains recovered from any extraintestinal infection in humans or animals. Although many virulence factors are associated with the pathogenicity of this group, it is difficult to identify or classify ExPEC strains based on a specific group of virulence genes [11]. Some studies developed molecular virulence patterns that define strains that harbor intrinsic extraintestinal virulence potential [12] or are potentially capable of causing urinary tract infection [13]. These molecular patterns are useful tools to track ExPEC both in the gastrointestinal tract or environment (soil, water, food), enabling the search for ExPEC reservoirs. Even though such methods may recognize the most virulent strains, they fail in identifying a considerable part of isolates recovered from clinical samples [13, 14]. The reason is that infections take place as a result of an imbalance between the virulence potential of the pathogen and the immune defenses of the host what makes it sometimes unclear whether the infection is being caused by a true pathogen or by an opportunistic strain. Considering this, the use of epidemiologic data and multi-locus sequence typing (MLST) for the identification of strains belonging to major pathogenic clonal groups could help in the determination of the potential pathogenic role played by an *E. coli* strain [15–17].

The emergence of MDR *E. coli* strains calls attention to the spread of clones carrying virulence along with resistance-encoding genes, making the control of these pathogens potentially difficult [2,18–22]. In this context, ST131 is the MDR high-risk clonal group widely disseminated and studied worldwide nowadays. Other clonal groups, presenting MDR phenotype, like ST405, ST38, and ST648, have also emerged and are already considered as of global risk [23]. On the other hand, some STs presenting MDR phenotype, although being isolated around the world, have not their pathogenic potential determined yet. The spread of MDR pathogens is a major Public Health concern that needs to be adequately addressed towards efficient control. Based on that, the World Health Organization (WHO) called attention to this problem and the need for alternative therapeutic options for the treatment of MDR infections. The development of vaccines and anti-virulence compounds could be alternative approaches to combat MDR strains, especially those showing pan drug resistance (PDR) phenotype [24]. However, for these alternatives to be effective, advanced knowledge is necessary, since not all pathogenic strains share the same virulence factors, and the use of the most prevalent virulence factors as targets can be problematic as they can adversely affect the gut microbiota.

It is well accepted that the phylogenetic grouping of *E. coli* keeps a very good correlation with the virulence potential of bacterial isolates. More recent epidemiological data have shown that non-virulent strains are mostly classified in the phylogenetic groups A, while diarrheagenic *E. coli* (DEC) are B1, and ExPEC are mainly B2. However, the fact that most *E. coli* virulence factors are carried on mobile genetic elements (e.g. plasmids and pathogenicity islands) may eventually cross these phylogenetic boundaries and promote the appearance of potential pathogens in atypical phylogroups. Thus, a global analysis of virulence and resistance characteristics of ExPEC isolates, especially those escaping typical classification, is essential for the understanding of the infections they cause as well as for devising therapeutic alternatives to cope with MDR *E. coli* strains. Aiming to provide information about the virulence determinants in non-typical ExPEC strains, we performed an extensive genotypic and phenotypic characterization of the MDR *E. coli* EC121 strain, which was characterized as belonging to phylogenetic group B1 and carrying only a few known virulence factors.

## Material and Methods

### Bacterial strain

The *E. coli* strain EC121 was isolated from the blood of a patient diagnosed with T-zone lymphoma and persistent infectious gastroenteritis, who had been hospitalized in a tertiary hospital located in the city of São Paulo, Brazil, in 2007. The patient died due to septic shock two days after the isolation of the EC121. The EC121 strain was kept frozen in glycerol at −80°C in ENTEROBACTERIALES-EXTRAINTESTINAL-EPM-DMIP collection n° A27A7C3. The initial virulence and resistance characterization showed that EC121 strain belonged to phylogroup B1, presented an MDR phenotype by routine susceptibility testing, and harbored few known virulence genes (*fim, hra, cvaC, ompA, ompT, sitA, iroN*). Furthermore, it was not considered a pathogenic strain because it harbored none of the virulence factors commonly involved in the characterization of ExPEC (presence of two of the following genes: *papA/C*, *sfaDE*, *afaBC*, *iuc/iut*, and *kpsMT*II) [25].

### Total DNA extraction, whole-genome sequencing (WGS), and genome assembly

The total bacterial DNA extraction was done using Wizard^®^ Genomic DNA Purification Kit (Promega - USA) following the manufacturer’s protocol. The extracted DNA was sequenced in an Illumina^®^ Hiseq1500 (Illumina-USA), using the Rapid protocol to obtain 2×250 paired-end reads, according to the manufactureŕs recommendations. Raw data were processed with Trimmomatic, and then the paired-end reads were assembled using SPAdes (version 3.12.0), with default parameters, and careful mode on [26].

### Genomic analyses and annotation

The obtained draft genome was submitted to various online bioinformatics platforms of the Center of Genomic Epidemiology (CGE) pipeline to determine (i) sequence types [27] for both *E. coli* MLST schemes; (ii) serotype (SerotypeFinder 2.0) [28]; (iii) presence and types of plasmid replicons (PlasmidFinder 2.0) [29]; (iv) presence of resistance genes (ResFinder 3.1) [30]; and (v) STEC virulence factors (VirulenceFinder 2.0) [31]. PHASTER [32] and PHAST [33] were used to detect bacteriophage sequences in each contig of the draft genome. The genome was annotated using Pathosystems Resource Integration Center (PATRIC) Comprehensive Genome Analysis service that uses RASTtk [34]. Each sequence that was assigned as a virulence factor in PATRIC’s database was manually submitted to BLAST/NCBI [35] and UniProt [36] to validate the virulence factors, to get all information about the virulence genes detected, to evaluate the completeness of the sequence and to determine its homology in relation to the RefSeq protein in Swiss-Prot. PATRIC [34] service was also used to build a phylogenetic tree using RAxML-VI-HPC or Fast tree 2, where all representative *E. coli* genomes from different pathotypes were used to construct the tree, as well as the deposited genomes of *E. coli* strains belonging to ST101 complex from diverse sources. The tree was built based on the concatenated sequence of all shared proteins among all genomes using RAxML or FastTree2. To construct the phylogenetic tree of EC121 and representative *E. coli* pathotypes, two *Escherichia fergusonni* strains ATCC35469 and NCTC12128 were used as outgroups. To build the phylogenetic trees, a total of 114 public genomes were randomly selected among the published genomes from the ST101 complex (ST101, ST359, ST2480, ST5957, and ST6388) using the PATRIĆs Genomes search tool. The *E. fergusonni* strains ATCC35469, *E. coli* str IAI1, *E. coli* O157:H7 str Sakai, *E. coli* O104:H4 str 2011c-3493 were used as outgroup. All phylogenetic tree final layout and annotation were done using iTOL v.4 [37]. The annotated genome was submitted to MacSyFinder from Galaxy@Pasteur [38] to detect CAS-CRISPR sequence type and the presence of secretion systems [39, 40].

### Serum agglutination assay for typing the O and H antigens

Serum agglutination assay was carried out following the standard methodology as described by Orskov and Orskov [41] for serotyping, using O serum against O100 and O154, and H serum against H25 provided by the Centers for Disease Control and Prevention (CDC, USA).

### Plasmid DNA extraction and analysis

Bacteria were cultivated in Tryptic soy broth (TSB - Difco, USA) at 37°C, in a static stove for approximately 18 h, and 1 mL of the culture was submitted to plasmid alkaline extraction protocol [42]. The *E. coli* strain 39R861 was used as a plasmid mass reference ladder and as control of extraction [43]. The plasmid extract was submitted to electrophoresis in an agarose gel (0.8%) in Tris-Borate-EDTA (TBE) buffer, stained with ethidium bromide solution (5 µg/mL), analyzed using Molecular Imager^®^Gel Doc™ XR^+^ with Image Lab™ Software System from Bio-Rad (USA). The molecular weight of each plasmid was calculated as previously described [44], based on migration in agarose gel of five different extraction assays followed by electrophoresis.

### *In silico* plasmid analysis

To accomplish the plasmid analysis, the following strategies were used. First, the draft genome was submitted to CGE to identify the contigs that contained replicons; subsequently, the assembled genome was analyzed by PlasFlow [45] to classify the possible source of each contig (as chromosomal or plasmid). The contigs containing replicons were analyzed using the Standard Nucleotide BLAST in NCBI.

### Data availability

The EC121 Whole Genome Shotgun project has been deposited at DDBJ/ENA/GenBank under the accession VYQD00000000. The version described in this paper is version VYQD01000000.

### Antimicrobial susceptibility testing

The minimum inhibitory concentration (MIC) was determined using the broth microdilution method, following the European Committee on Antimicrobial Susceptibility Testing (EUCAST) recommendations and breakpoints [46]. The following antimicrobials (Sigma - USA) were tested: ampicillin, piperacillin/tazobactam, ceftriaxone, ceftazidime, cefepime, aztreonam, ertapenem, imipenem, meropenem, ciprofloxacin, amikacin, gentamicin, tigecycline, colistin, polymyxin B, trimethoprim/sulfamethoxazole, and chloramphenicol. *E. coli* ATCC 25922 and *Pseudomonas aeruginosa* ATCC 27853 were used as quality control strains.

### Conjugation assay

Conjugation assay was conducted to resolve the EC121 conjugative plasmids. In the mating pair, strain EC121 was the donor strain whereas *E. coli* K-12 derived strains, J53 [47] and C600 [48], resistant to sodium azide, were the recipient strains. Conjugation was performed using overnight cultures of the donor and the recipient strains grown in LB, in the proportion of 1:2, respectively. One milliliter of fresh LB was added to the mating mixtures, following 3 hours incubation at 37 °C under static condition. After the incubation period, 100 µl of each mating mixture was plated into selective MacConkey agar plates supplemented with three different antibiotic combinations (20 µg/mL gentamycin and 100 µg/mL sodium azide; 2 µg/mL cefotaxime and 100 µg/mL sodium azide; and 30 µg/mL chloramphenicol and 100 µg/mL sodium azide). The colonies that grew in the selective medium (Transconjugants) were purified in the same selective medium and then stored for further characterization.

### Characterization of conjugative plasmids

The transconjugants obtained were analyzed by PCR for the presence of virulence encoding-genes, resistance *bla*_CTX-M-2_ gene, and determination of the replicon types. Additionally, the susceptibility profile of the transconjugants was determined by the minimum inhibitory concentration (MIC) method [14,49,50].

Determination of the lowest bacterial inoculum which was resistant to human **serum complement.**

To access the bacterial serum-resistance, the lowest bacterial inoculum resistant to serum was assessed. Lyophilized human complement serum (Sigma, USA) was reconstituted in sterile phosphate-buffered saline (PBS). The assay was performed in 96-wells plates, where complement serum was distributed in each well (90 uL per well). Bacteria were grown overnight at 37°C, serially diluted (1:10) in complement serum until 10^-10^ and incubated at 37°C. Aliquots of 10 µL of each well were seeded onto MacConkey agar plates after 30 min, 1 h and 2 h of incubation. Simultaneously, another assay was performed with previously heat-inactivated serum as control. The *E. coli* strains J96 and C600 were used as resistant and susceptible controls, respectively [51]. The lowest bacterial inoculum resistant to human complement was determined by the last bacterial dilution which had bacterial growth onto MacConkey after the challenge. For each assay, the initial bacterial inoculum was determined by diluting bacteria in PBS, plating in MacConkey agar and CFU counting. The data was reported in CFU/mL. Biological assays were performed in triplicates.

### Biofilm formation on abiotic surfaces

Biofilm formation was evaluated on polystyrene and glass surfaces as described by Lima et al [49] in a 24 hour-assay using the following media: DMEM high-glucose and TSB. Each assay was performed in biological and experimental triplicates. The EAEC 042 and laboratory *E. coli* HB101 strains were used as positive and negative controls, respectively; in all assays, a non-inoculated well was used as control of dye retention, and the prototype strain CFT073 as an ExPEC control.

### Cell culture and maintenance

HeLa (ATCC^®^ CCL-2^™^), intestinal Caco-2 (ATCC^®^ HTB-37^™^) and bladder T24 (ATCC^®^ HTB-4^™^) cell lineages were used to evaluate the ability of strain EC121 to interact with eukaryotic cells. HeLa and Caco-2 cells were cultured in DMEM, High Glucose, GlutaMax™ (Gibco-ThermoFisher Scientific, USA), supplemented with 10% bovine fetal serum (BFS) (Gibco, USA), 1% non-essential amino acids (Gibco, USA), and 1x Penicillin-Streptomycin-Neomycin (PSN) antibiotic mixture (Gibco, USA), while T24 cells (ATCC HTB-4) were cultured in McCoy 5A (modified) media (Gibco, USA), supplemented with 10% of BFS and 1x PSN antibiotic mixture. All lineages were kept at 37°C in an atmosphere of 5% CO_2_. For all assays, cell suspensions containing 10^5^ cells/mL were seeded in 24-well plates, with or without glass coverslips for qualitative or quantitative assays and cultured from two up to 10 days until.

### Adherence assay in HeLa, Caco-2, and T24 cells

The adherence properties of EC121 were evaluated qualitatively and quantitatively. In both assays, the epithelial cells were washed three times with PBS and 1 mL of proper media, supplemented with 2% of BFS, was added. After the bacterial inoculation, the qualitative and quantitative assays were incubated for 3 hours at 37 °C, then washed three times with PBS and processed according to the type of the test. The qualitative assays were performed with all cell lineages (HeLa, T24, and Caco-2) using 20 of inoculum, obtained from standardized bacterial cultures grown overnight, and after the incubation and washes, were fixed and stained as described previously [52]. The quantitative assay was carried out using HeLa cells in full confluency to evaluate the efficiency of the EC121 adherence in the presence and absence of 2% of D-mannose. The multiplicity of infection (MOI) of 50, obtained from an overnight culture, washed, and adjusted in PBS, confirmed by O.D. and bacterial count, was used as inoculum. After incubation and washes, the cells were lysed by adding 1 mL of sterile bi-distilled water, which was recovered, diluted, and plated onto MacConkey agar for quantification as previously described [53]. The assays were performed in biological duplicates and experimental triplicates, the data were expressed as SEM. The *E. coli* strains C1845, CFT073, and C600 were used as controls.

### Short period interaction and invasion assay in Caco-2, and T24 cells

The invasion assays were carried out using Caco-2 and T24 cells in full confluency, as described by Martinez et al. [54], with modifications, in two sets of plates simultaneously. The infection was done using a MOI of 50, and the assays were incubated for 2 h, at 37°C, in a normal atmosphere. After this period, one plate set was washed three times with PBS and incubated again with PBS containing 100 µg/mL of amikacin for 1 hour at 37°C, to kill all extracellular bacteria. After the incubation period, the assay was washed three times to remove all antibiotics, cells were lysed with water, and the well contents were collected, diluted and plated onto MacConkey agar to obtain the number of internalized bacteria. The other set was washed with PBS three times, the cells were lysed, and contents of each well were collected, diluted and plated to obtain the total number of bacteria interacting with the cells in the period. An aliquot of the PBS recovered from the last wash after incubation with amikacin was collected and plated without dilution, to ensure that the treatment had killed all extracellular bacteria. The total interaction value was the percentage of the total bacterial associated with the cells in relation to the initial inoculum. The invasiveness was determined by the ratio between the number of internalized bacteria and the initial inoculum, expressed in percentage. The *Escherichia albertii* strain 1551::*eae* was used as adherent and non-invasive control [55], the *E. coli* strain C600 as a negative control, and the CFT073 as an ExPEC control. The assays were performed in biological and experimental triplicates, and data were reported as SEM.

### In vivo assay in Galleria mellonella virulence model

The full virulence potential of EC121 was evaluated using the virulence model of *G. mellonella* as previously described [56, 57]. Briefly, 10 mL of logarithmic phase cultures were washed twice and adjusted to the optical density of 0.7 (O.D.595) in 0.85% NaCl and serially diluted to obtain a bacterial concentration of 1×10^6^ CFU/mL. Volumes of 10 µl of the bacterial suspension were injected in the left larvae proleg using a Hamilton syringe (26S gauge, 50 µL capacity). The tests were performed in three independent assays, each of them using 5 larvae per bacterial strain tested. The *E. coli* strains CFT073 and C600 were used as positive and negative controls, respectively, and saline injection as the procedure control.

### Statistical analyses

Student t-test was applied to calculate statistical significance. For biofilm production, the Wilcoxon matched-pairs test correction was used, and for the cell interaction and invasion assays, the Wilcoxon-Mann-Whitney test was applied. The Kaplan-Meier survival curve was done for survival analysis, and the difference between the groups was determined by Log-rank (Mantel-cox) test and Gehan-Breslow-Wilcoxon test. The threshold for statistical significance was a *P*-value < 0.05. The analyses were performed in Prism 5.0 (GraphPad Prism Software, Inc).

## Results

### Genetic characterization and *in silic*o analysis

The genome assembly of the strain EC121 generated 143 contigs, with a total genome predicted size of 5,119,556 bp, and 50.43% of GC content. PlasFlow algorithm identified 43 of 143 contigs as chromosomal and other 66 contigs as plasmidial. The contigs assigned as plasmidial comprised at least 426,419 bp of the genome.

### EC121 belongs to serotype O154:H25, ST101-B1, and is related to diarrheagenic *E.* ***coli*.**

The MLST analyses showed that the EC121 strain belonged to the ST101/ST88, according to the Warwick and Pasteur MLST schemes, respectively. The determination of serotype type by sequencing analysis was inconclusive, because two possible serotypes, O154:H25 and O100:H25, could be assigned. Using specific serum agglutination assays it was determined that strain EC121 expressed the O154 antigen. Thus, it was characterized as belonging to serotype O154:H25. Moreover, group IV capsule encoding genes were identified on its genome. A phylogenetic tree built using reference *E. coli* strains from all pathotypes showed that EC121 was related to diarrheagenic *E. coli* (DEC) strains, since it was positioned in a clade closely associated with Shiga toxin-producing *E. coli* (STEC) (Figure 1).

**Figure 1.**
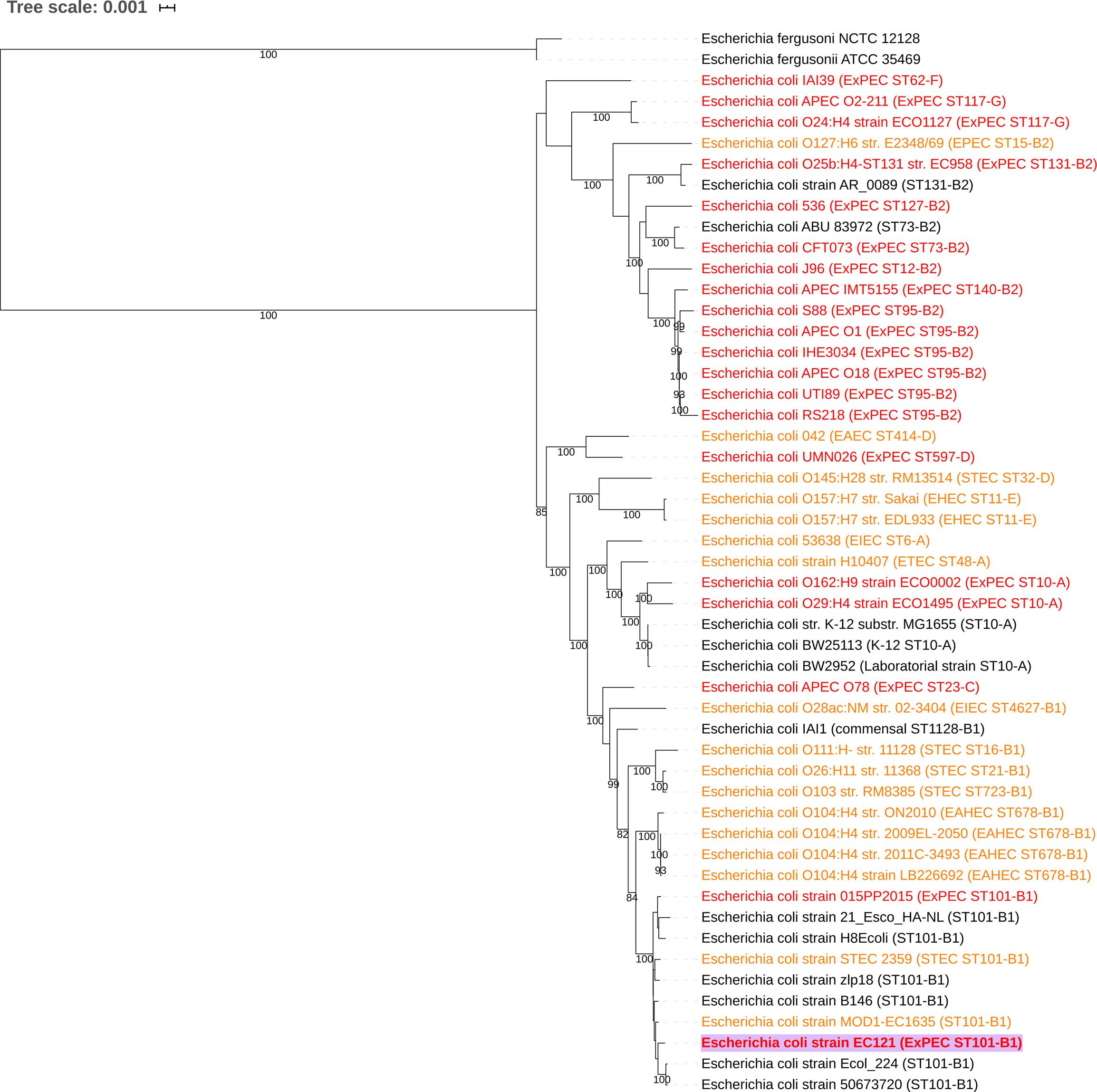
EC121 Phylogenetic tree. A phylogenetic tree was built with genomes of reference *E. coli* strains, of relevant pathogenic strains from all *E. coli* pathotypes, and some strains from ST101, using Maximum Likelihood-based algorithm (RAxML) in PATRIC. In parenthesis, the strain pathotype (when known), the Sequence type, and phylogroup according to ClermonTyping [105] are depicted. Diarrheagenic *E. coli* strains were in orange; Extraintestinal pathogenic *E. coli* strains were in red, commensal or strains that the origin was not described were in black. In bold and with purple label background, the strain studied in present work. Bootstrap upper than 80 were informed in the tree.

A second phylogenetic tree was built with 95 strains available at the NCBI, which belonged to the ST101 complex and were recovered from distinct sources (Figure 2, Table S1). Analysis of the tree showed that the majority of the strains of this ST are MDR, many of them carrying *mcr*-1 (mobile colistin resistance gene), a variety of β-lactamases (*bla*_CTX-M_-like, *bla*_OXA_-like, *bla*_NDM_-like) and genes related to fosfomycin resistance (*fosA3*). Interestingly, these strains were isolated from food, environment, animals, and humans, as part of the microbiota or involved in both intestinal and extraintestinal infections, in all continents. Among the strains isolated from human infections (Figure S1 and Table S1), most were diagnosed as extraintestinal pathogens (28 strains), while three were intestinal pathogens; among the latter, one was Shiga toxin-producing *E. coli* (STEC), and one was enterotoxigenic *E. coli* (ETEC). Regarding the isolates from food and animals, seven strains were identified as STEC (Figure 2, Figure S1 and Table S1). Showing that ST101 is associated with intestinal and extraintestinal infection and it is a clonal group associated with MDR phenotype. The full resistome and isolation data (accession number, year and country of isolation, etc.), related to strains from the ST101 complex used to build the phylogenetic trees, were provided in Table S.

**Figure 2.**
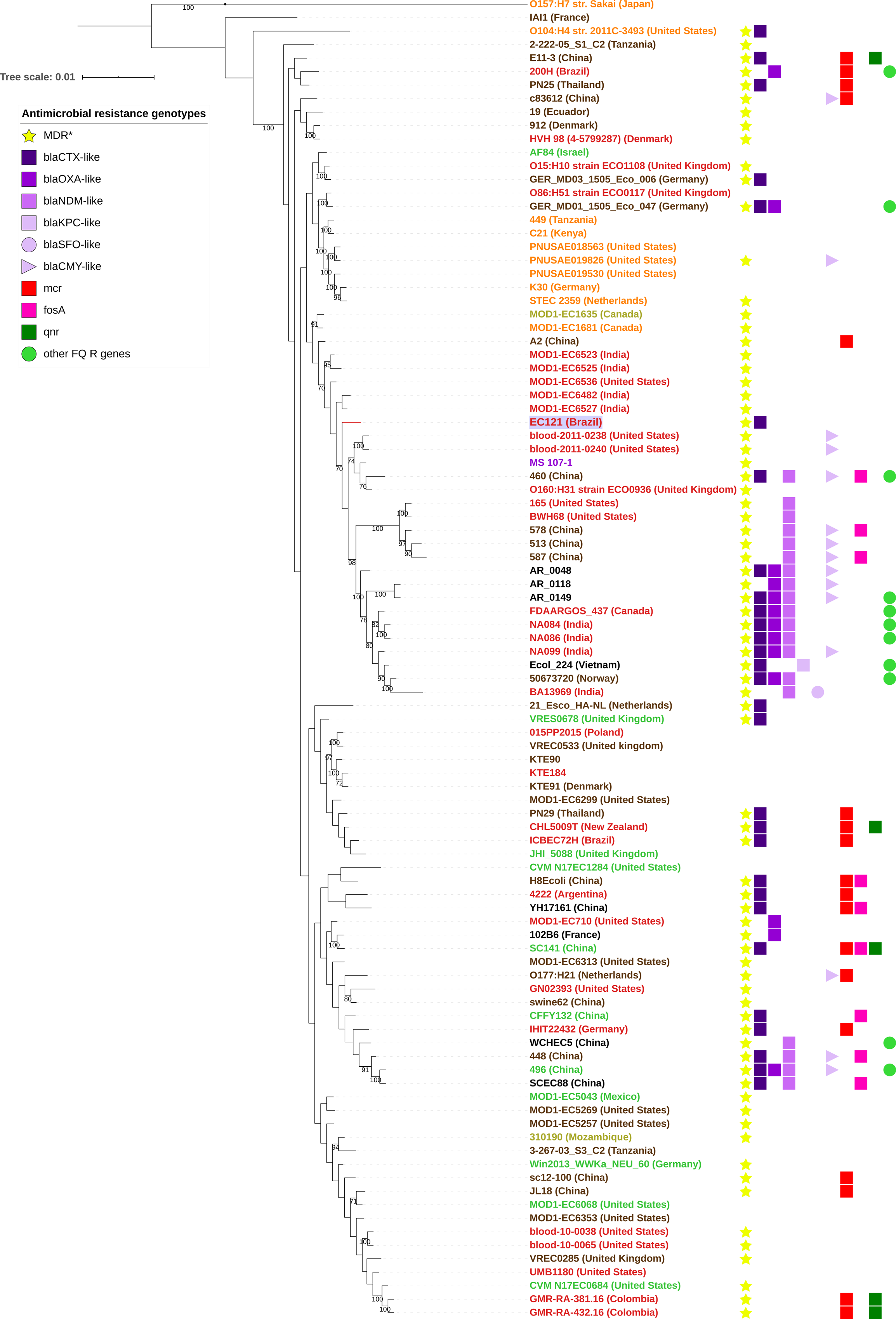
ST101 complex phylogenetic tree. The phylogenetic tree was built using 95 *E. coli* strains from ST101 complex (ST101, ST359, ST2480, ST5957, and ST6388) from diverse source and countries using all shared protein among them on FastTree2 to build the tree. *E. coli* strain IAI1, *E. coli* O157:H7 str Sakai, and *E. coli* O104:H4 str 2011c-3493 are used as outgroups. Bootstrap upper than 50 were informed in the tree. Label colors are related with isolates origin or host diseases, being Shiga toxin-producing *E. coli* (STEC) in orange independently of the origin; Diarrheagenic *E. coli* (DEC) strains in yellow; Extraintestinal pathogenic *E. coli* (ExPEC) strains in red; isolates from microbiota in brown; isolates from retail food or environment in green; strains that the origin was not described in black. One strain isolated from Crohn’s disease is in purple. In bold and with purple label background, the strain of present work. * All strains that harbor AMR for 3 or more antimicrobial classes were designed as MDR. Other FQ (Fluoroquinolone) resistance genes detected were the mobile genes *qepA* and *aac(6’)-Ib-cr*. Mutations that confer resistance to FQ were not considered to build the AMR information in this tree.

**Table 1.** EC121 annotation overview.*^a^*

| Protein features | Occurrence |
| --- | --- |
| Hypothetical proteins | 702 |
| Proteins with functional assignments | 4,473 |
| Proteins with E.C.<sup>b</sup> number assignments | 1,300 |
| Proteins with G.O.<sup>b</sup> assignments | 1,076 |
| Proteins with Pathway assignments | 911 |
| Proteins with PLfam<sup>b</sup> assignments | 5,063 |
| Proteins with PGfam<sup>b</sup> assignments | 5,064 |

*a.* Results obtained using PATRIC annotation service;
*b.* Abbreviations: E.C. number - Enzyme commission universal G.O. - Gene Ontology
Consortium; PLfam- PATRIC genus-specific family; PGfam- PATRIC cross-genus
family.

### EC121 harbors genes involved in virulence and stress response

The EC121 genome annotation showed that strain EC121 contained 5,175 coding sequences (CDS), 82 tRNA, and 13 rRNA. One CRISPR *locus* was identified as type 1-IE and presented two arrays and 30 CRISPR-repeat regions (Figure 3A). Among the CDSs annotated, 702 corresponded to putative proteins designated as hypothetical proteins, and 4,473 CDS to putative proteins with functional assignments. Of interest, 221 genes were reported as belonging to systems involved in response to stress, virulence, and defense (Table 1 and Figures 3B-C). Analysis using the MacSyFinder tool identified 10 type V secretion system proteins, nine of which were from type T5aSS and one from T5cSS (EhaG) (Table S2). One type III secretion system similar to *Salmonella* T3SS, and an incomplete type VI secretion system that carried only *tssB*, *tssD*, *tssE*, *tssH*, and *tssI* genes were also present (Table S2). All genes that were reported by the PATRIC virulence factor database were manually curated to provide information about their full sequence. As shown in Tables 2, S3, and S4, the genome of the EC121 strain encodes multiple adhesins, invasins, iron uptake systems, and genes involved with the evasion of the host immune system. By the position of strain EC121 in the pathotype phylogenetic tree, some of these virulence factors are related to the pathogenesis of DEC, specifically of STEC and ETEC (Tables 2 and S3). Other virulence factors found were related to *Salmonella* spp. (PagN adhesin and systems associated with immune evasion and macrophage survival) and *Shigella* spp. (genes associated with intracellular survival and spread). Moreover, many accessory genetic clusters associated with the bacterial ability to cause extraintestinal infections, i.e., genes involved with biofilm formation, adherence to extraintestinal cells, iron acquisition and immune evasion were also detected in the EC121 genome. Furthermore, other clusters associated with Urinary Tract Infections (UTI) were detected (Tables 2 and S3).

**Figure 3.**
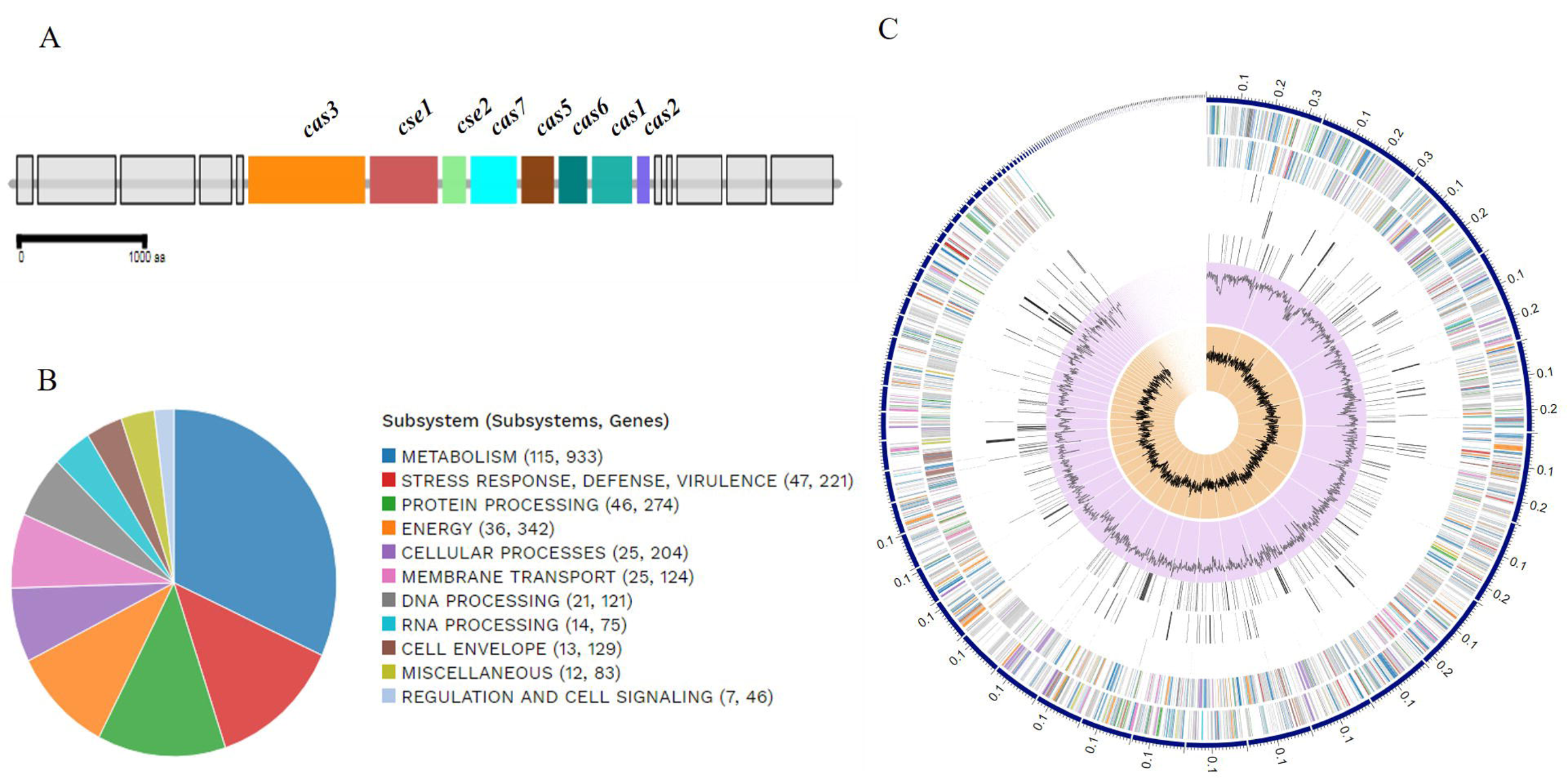
CRISPR locus composition and biological systems assignment in EC121**. A –** Genomic architecture representation of CRISPR *locus* in the EC121 genome; the image was obtained in MacSyFinder from genome annotation; **B -** Representation of EC121 genome composition in subsystems based on protein biological data obtained *in silico*; **C -** EC121 genome schematic composition, based on annotation, ordered by contig size. In circle from outer to the inner, forward strand, reverse strand, RNA related genes, antimicrobial resistance, virulence factors, GC content, and GC skew. The colors in forward and reverse strands correspond to the subsystems presented in B. Figures 2B and 2C were obtained using the comprehensive genome analysis service at PATRIC.

**Table 2.** Complete virulence factors identified

| Virulence traits | Associated with virulence in |  |  |  |
| --- | --- | --- | --- | --- |
|  | Other Genera | DEC | ExPEC | Various |
| **Colonization and invasion** | MlaA, MsbB, Sfm, MisL, IcsP | Lpf-1<sub>O26</sub>, EhaG, Hcp, Elf, EhaA, EhaB, CFA-I | FdeC, Pix, Ygi, Yad, Yeh, Yra, Yfc, YchO, IbeB, IbeC, EptC, OmpA | Type 1 Fimbriae (H190), Ecp, Curli |
| **Immune evasion** | SodB, TrxA, SirA, FpkA |  | Iss, OmpTp<sup>a</sup>, OmpTc<sup>a</sup>, Mig-14, HlyF | RelA |
| **Iron acquisition** |  |  |  | Sit, Iro, Fhu, Enterobactin |
| **Regulators** | DsbA, DegP, SlyA, CpxAR | EvgAS | DsbAB, PhoPQ | QseBC, RcsAB |
| **Toxins and Bacteriocin** |  |  | Microcin V, Colicin B, Colicin M | ClyA, Hly III |
| **Non-LEE effectors** |  | EspL1, EspL4, EspX1, EspX4, EspX5, EspR1 |  |  |

<sup>a</sup> OmpTc for chromosomal variant of OmpT protein, and OmpTp for plasmid variant of OmpT protein.

### No complete phage sequences were detected in the EC121 strain

**The search for** phage sequences in strain EC121 identified 12 regions containing genes from a variety of different phages, ranging from 6 to 31 kb (Table 3, Table S4). Although the database considered a predicted phage sequence in region 6 as intact, based on their score criteria, it’s size (14,400 bp) was not compatible with the size of the predicted “*Salmonella* phage Fels-2”, whose complete genome sequence deposited in NCBI database is 33,693 bp. Remarkably, parts of the cytolethal distending toxin (Cdt-I and Cdt-V), and Shiga-like toxin (Stx1a and Stx2c) converting phages, as well as of *Shigella* serotype-converting phages SfI, SfII, and SfV were detected among these regions (Table 3 and Table S4). However, no toxin-encoding genes were identified in strain EC121.

**Table 3.** Predicted phages detected in the EC121 genome.

| **Region** | **Length** | **Completeness<sup>a</sup>** | **Most Common Phage/accession number** | **GC %** |
| --- | --- | --- | --- | --- |
| **1** | 6.3 kb | Incomplete | *Bacillus* phage G/NC_023719 | 51.27 |
| **2** | 30.9 kb | Incomplete | *Salmonella* phage 118970_sal3/NC_031940 | 49.82 |
| **3** | 14.2 kb | Questionable | Enterobacteria phage P88/NC_026014 | 49.75 |
| **4** | 20 kb | Incomplete | *Shigella* phage POCJ13/NC_025434 | 47.95 |
| **5** | 28.5 kb | Incomplete | *Shigella* phage SfII/NC_021857 | 45.37 |
| **6** | 14.4 kb | Intact | Enterobacteria phage Fels-2/NC_010463 | 49.56 |
| **7** | 9.4 kb | Incomplete | *Bacillus* phage Shanette/NC_028983 | 49.50 |
| **8** | 7.5 kb | Incomplete | Enterobacteria phage phi92/NC_023693 | 44.44 |
| **9** | 10.2 kb | Questionable | Enterobacteria phage P2/NC_001895 | 55.39 |
| **10** | 3.8 kb | Incomplete | Bacteriophage WPhi/NC_005056 | 50.16 |
| **11** | 8.4 kb | Questionable | Phage cdtI/NC_009514 | 46.05 |
| **12** | 6.7 kb | Incomplete | *Shewanella* sp. phage 1/4/NC_025436 | 47.92 |

*a.* Phage completeness was determined by scores obtained in the search algorithm (PHASTER) based on the number of the phage's specific CDSs detected in the analyzed region.

**Table 4.**
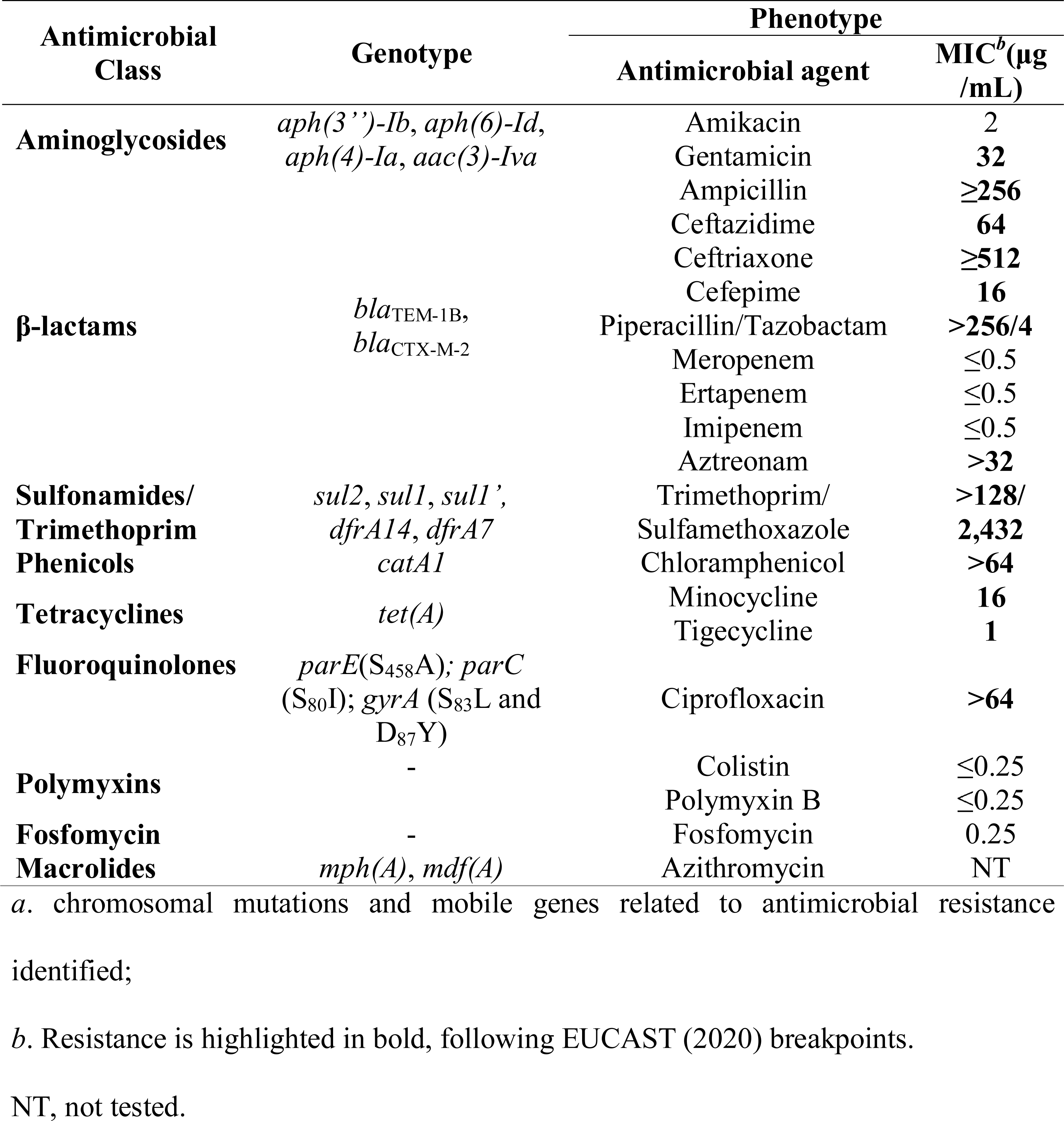
Antimicrobial resistance genotype and phenotype observed in strain EC121.

### EC121 has multiple resistance genes, and several efflux-pumps compatible with its **antimicrobial susceptibility profile.**

The software ResFinder identified 15 resistance genes in strain EC121, which are involved in reduced susceptibility to aminoglycosides, β-lactams, macrolides, phenicols, sulphonamides, trimethoprim, and tetracyclines (Table 4). Mutations in *parE*, *parC*, and in *gyrA* genes, which confer resistance to fluoroquinolones, were also observed in the EC121 genome. Additionally, several efflux-pumps related to resistance to heavy metals (copper and mercury), arsenic, disinfectants (QacE), and antimicrobials (AcrAB-TolC, AcrAD-TolC, AcrEF-TolC, AcrZ, EmrAB-TolC, EmrD, EmrKY-TolC, MacA, MacB, MdfA/Cmr, MdtABC-TolC, MdtEF-TolC, MdtL, MdtM, and SugE) were also detected. The MDR phenotype profile of strain EC121 assessed by the microdilution method was consistent with the genomic findings, as can be seen in Table 4.

### The EC121 strain harbors multiple plasmids

Four bands were detected by agarose gel electrophoresis, suggesting that EC121 harbor multiple plasmids (Figure S2). Seven different replicons were found in EC121 using PlasmidFinder 2.0 (IncHI2A, IncHI2, IncQ1, IncFII, IncFIB, IncN, and IncM1). Although the EC121 genome is still in draft, we analyzed the replicon containing contigs identified by the PlasmidFinder to provide more information about the strain’s plasmidial content. One contig of 128,478 bp in length bears both IncHI2A and IncHI2 replicons. The contig’s BLAST showed high identity with the pYps.F1 plasmid of the *Yersinia pseudotuberculosis* strain Yps.F1 (cover: 99%, e-value: 0.0, identity: 99.77%) (Figure S3). The IncM1 replicon was identified in one contig of 65,565 bp in length, which was circularized in the assembling, and showed high homology (100% of identity and coverage) with the pASM2 plasmid (accession n° NZ_CP019841.1) of *Enterobacter roggenkampii* strain R11 (Figure S4). The other replicons were segregated into four different contigs, with IncFIB and IncFII into contigs of 37,695 and 30,674 bp in length, respectively, and IncN and IncQ1 into contigs with less than 3.5 kb each. The IncFIB replicon’s contig also contained the virulence genes *iroN, iss,* and *traT* as identified in the sequence. The *in silico* analyzes suggested that the plasmid was similar to the pAPEC plasmid that carries virulence and resistance genes simultaneously. Although the IncFII replicon was identified into a different contig, the data indicated that both the IncFII and IncFIB replicons represent segments of the same plasmid. Further studies are required to unravel this plasmid genetic composition. Conjugation assays were performed to evaluate the presence of conjugative plasmid in EC121, to this purpose using agar plates with different antimicrobial combinations (sodium azide [100 µg/mL] with gentamycin [20 µg/mL] or cefotaxime [2 µg/mL] or chloramphenicol [30 µg/mL]) to select the transconjugant strains. While no chloramphenicol-resistant transconjugant colony was detected, five transconjugants were recovered from the gentamycin selective agar plate and 20 from the cefotaxime selective agar plate. Some colonies were purified and then investigated regarding their replicon type and presence of virulence genes that are generally located in plasmids (*hlyF, sitA,* and *iroN*) by PCR. All cefotaxime-resistant transconjugants carried the IncHI2 replicon, *bla*_CTX-M-2_ and lacked the virulence genes investigated, whilst all five gentamycin-resistant transconjugants carried multiple replicons (IncFIB, IncM1, IncN, and IncHI2), as well as the *hlyF, sitA*, and *iroN* genes (Table 5), and *bla*_CTX-M-2_ resistance gene. These results suggested that the *bla*_CTX-M-2_ gene was inserted into IncHI2A/IncHI2 plasmid.

**Table 5.** Resistance profile, replicon type, and virulence genes identified in the EC121’s plasmids transferred to *E. coli* K-12 strains

| Traits | EC121 | J53 | C600 | ACG04 | ACC09 |
| --- | --- | --- | --- | --- | --- |
| **Antimicrobial <sup>a</sup>** |  |  |  |  |  |
| Ceftazidime | 64 | 0.25 | 0.25 | **8** | **2** |
| Ceftriaxone | ≥64 | ≤0.125 | ≤0.125 | >**64** | >**64** |
| Cefepime | 16 | 0.25 | ≤0.125 | **16** | **2** |
| Piperacillin/<br>Tazobactam | >256 | 2 | 2 | **8** | **2** |
| Aztreonam | >32 | ≤0.125 | ≤0.125 | **16** | **4** |
| Amikacin | 4 | 2 | 2 | **16** | **2** |
| Gentamycin | 16 | 0.5 | 0.5 | **32** | 0.5 |
| Trimethoprim/<br>Sulfamethoxazole | >128 | ≤0.25 | ≤0.25 | >**128** | >**128** |
| Ciprofloxacin | >64 | ≤0.125 | ≤0.125 | ≤0.125 | ≤0.125 |
| Minocycline | 16 | 1 | 1 | **4** | 1 |
| Tigecycline | 1 | 0.25 | 0.125 | 0.25 | 0.125 |
| **Replicon type <sup>b</sup>** |  |  |  |  |  |
| IncHI2 | + | - | - | + | + |
| IncL/M | + | - | - | + | - |
| IncFIB | + | - | - | + | - |
| IncN | + | - | - | + | - |
| **Virulence <sup>c</sup>** |  |  |  |  |  |
| *hlyF* | + | - | - | + | - |
| *iroN* | + | - | - | + | - |
| *sitA* | + | - | - | + | - |
| **Resistance gene** |  |  |  |  |  |
| *bla<sub>CTX-M</sub>* | + | - | - | + | + |

<sup>a</sup> The antimicrobial resistance phenotype of transconjugant strains was assessed by minimum inhibitory concentration (MIC), performed using the broth microdilution method according to EUCAST guideline, and was expressed in µg/mL. Bold values indicate an increased MIC value when comparing strains bearing plasmids with its receptor without plasmid.
<sup>b</sup> Replicons identified in EC121 by PlasmidFinder, and screened by PCR;
<sup>c</sup> Virulence genes identified in EC121 genome and screened by PCR;
+ for presence of the gene; - for the absence of the gene;

**Table 6.** Estimated bacterial inoculum resistant to serum activity (CFU/mL).

| Challenge period | Strains |  |  |  |  |  |
| --- | --- | --- | --- | --- | --- | --- |
|  | J96 |  | EC121 |  | C600 |  |
|  | NHS | inHS | NHS | inHS | NHS | inHS |
| 30 min | 10<sup>1</sup> | 10<sup>1</sup> | 10<sup>2</sup> | 10<sup>1</sup> | 10<sup>8</sup> | 10<sup>1</sup> |
| 1 h | 10<sup>2</sup> | 10<sup>1</sup> | 10<sup>2</sup> | 10<sup>1</sup> | NG | 10<sup>1</sup> |
| 2 h | 10<sup>2</sup> | 10<sup>1</sup> | 10<sup>2</sup> | 10<sup>1</sup> | NG | 10<sup>1</sup> |

*a.* Values represent the approximate relative mean of the lowest bacterial inoculum that remained viable after the challenge. All assays were performed in triplicate using 50% serum diluted in PBS (v/v); NHS - Normal human serum; inHS - inactivated human serum; NG - no growth after the challenge.

One transconjugant strain of each type was submitted to the microdilution assay to verify the antimicrobial resistance phenotype. The *E. coli* strain ACC09 (*E. coli* J53 harboring the IncHI2/IncHI2A plasmid) obtained from the cefotaxime agar plate presented a resistant phenotype to ceftriaxone and co-trimoxazole and reduced susceptibility to all beta-lactams (Table 5), whereas the *E. coli* strain ACG04 (*E. coli* C600 carrying multiple plasmids) had reduced susceptibility to all the antimicrobial tested except for ciprofloxacin and tigecycline, and presented increased MIC values to all beta-lactams when compared to ACC09 (Table 5). Although we have isolated only the IncHI2/IncHI2A plasmid, multiple plasmids were transferred efficiently to the receptor strain in a short period of conjugation (3 hours) indicating that they were all conjugative or mobilizable plasmids.

### Virulence phenotype

To evaluate the expression of the virulence-encoding genes detected, *in vitro* assays were performed to analyze the ability of the strain to: (i) resist to the bactericidal activity of the serum complement system; (ii) attach to abiotic surfaces and form biofilms; (iii) adhere to and invade eukaryotic cells. Besides, *in vivo* assays were performed using the *G. mellonella* infection model to evaluate the EC121 virulence potential.

### The EC121 strain was resistant to the bactericidal activity of the complement **system and adhere to abiotic surfaces.**

To disseminate in the host, extraintestinal pathogenic bacteria must be able to survive the serum bactericidal activity. To identify such a feature in EC121, we determined the lowest serum-resistant bacterial inoculum using a pool of normal human sera (NHS). The lowest inoculum of EC121 strain that resisted serum activity after two hours was 10^2^ CFU/mL, which was similar to the one obtained for the resistant control strain J96. *E. coli* strain C600, used as the susceptible control, barely resisted to 30 min-exposition period in the highest inoculum tested (10^8^ CFU/mL). To validate if the bacterial survival was associated with the resistance to complement activity, assays were repeated with heat-inactivated serum. In this condition, all strains survived the challenge with similar inoculum (Table 6). The ability to adhere to the abiotic surface and form biofilm can confer many advantages to any pathogen, including persistence in particular niches and tolerance against antimicrobials and the host immune system. The EC121 strain was able to adhere to borosilicate coverslips and polystyrene when grown in DMEM and TSB, as shown in Figure 4. Although its adherence to the abiotic surface was not massive as the adherence presented by the positive control strain (EAEC 042), it was significantly more intense than that of the negative control strain (HB101) (Figures 4). Furthermore, EC121 produced similar (*P* > 0.05) or higher (*P* <0.01) biofilm masses as the ExPEC prototype strain CFT073 in DMEM and TSB, respectively (Figure 4b).

**Figure 4.**
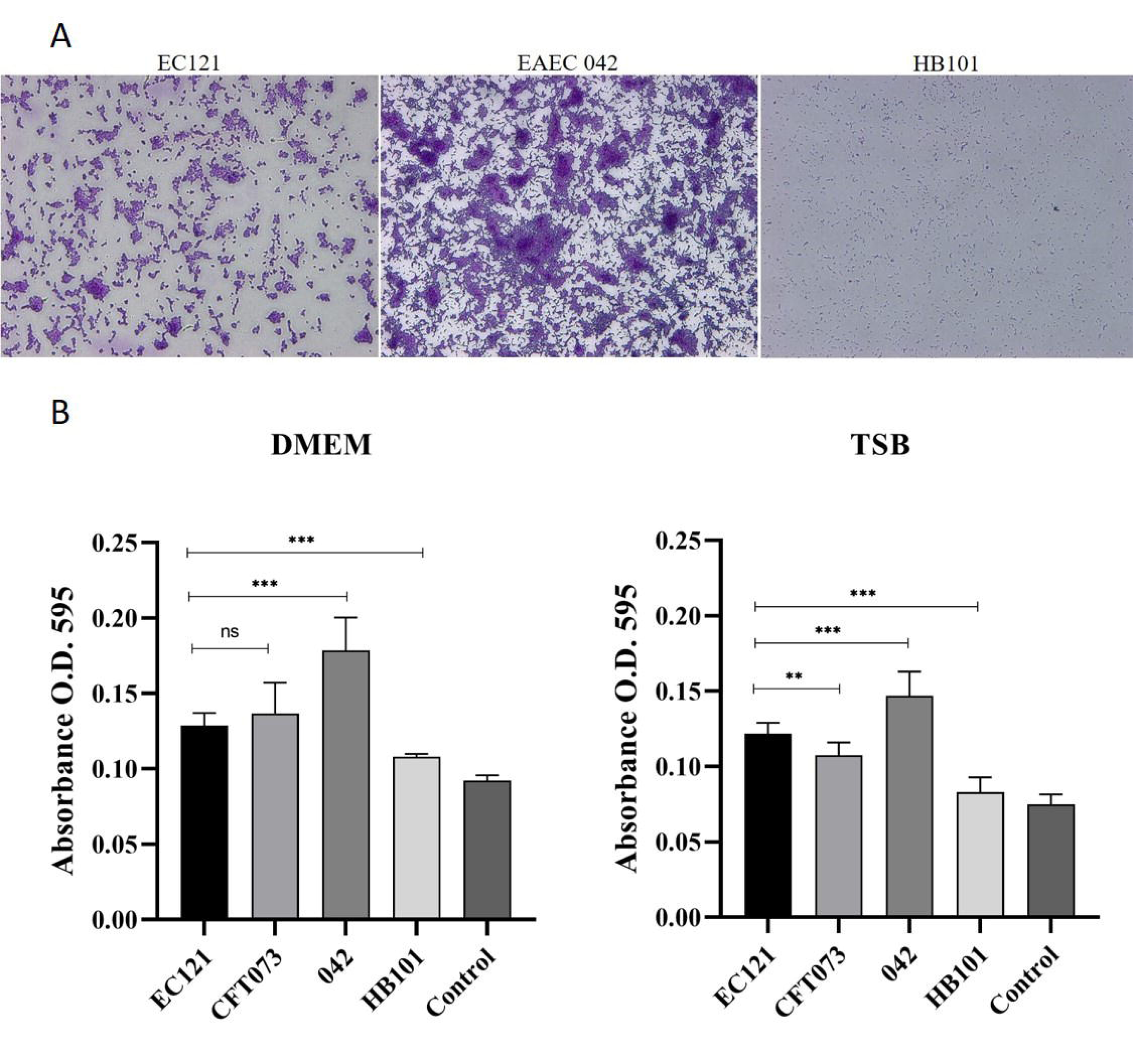
Biofilm formation on abiotic surfaces. A. Qualitative assay showing the EC121 adhered to glass coverslips after 24h of incubation at 37 °C, in DMEM, EAEC 042 and HB101 were used as positive and negative controls, respectively. Bacteria were stained with crystal violet, observed in optical microscopy (OM) 400 x **B.** Quantitative biofilm assays were performed in DMEM and TSB, at 37°C, for 24 h, comparing EC121 capacity to produce biofilm in polystyrene surface. EAEC 042 was used as positive control, CFT073 as an ExPEC control and HB101 as negative control. *P* values: ** *<*0.01; *** *<*0.001; ns >0.05.

### The EC121 strain adheres to and invades eukaryotic cell lineages

**At first, we** accessed, qualitatively, the ability of strain EC121 to adhere to HeLa, T24, and Caco-2 cells, using 3-h adherence assay without D-mannose. This assay showed that EC121 interact efficiently with all cells tested (Figure S5). Subsequently, a quantitative 3-h adherence assay in HeLa cells using initial inoculum of 1×10^7^ was performed in the absence of D-mannose, except for EC121, which also was tested in the presence of 2% of D-mannose. EC121 was able to adhere to, both in the presence or absence of D-mannose, which abolishes the adherence mediated by type-1 fimbriae. However, the presence of D-mannose reduced significantly the adherence ability of strain EC121, lowing the adherence in about 90% (Figure 5), from 1.3×10^7^ CFU/mL to 1.58×10^6^ CFU/mL (Figure 5). Noteworthy, EC121 adherence levels in the absence of D-mannose was higher than all controls tested in the same conditions (*P* < 0.01) (Figure 5). We also investigated whether the EC121 strain was able to invade T24 and Caco-2 cells. In a short-period invasion assay, the EC121 strain interacted with T24 cells as efficiently as CFT073 but invaded the cell lineage significantly more (P < 0.005) (Figure 6). On the other hand, its interaction with differentiated and polarized Caco-2 cells was lower when compared with CFT073, although there was no difference related to their capacity to invade this type of cell lineage on the conditions tested. The capacity to interact and invade eukaryotic cells in the presence of D-mannose was also assessed to evaluate whether the interaction and invasion abilities of EC121 were dependent on type-1 fimbriae or another mannose-dependent adhesin. The EC121 interaction with both lineages reduced significantly in the presence of D-mannose, like in HeLa cells showing that mannose-sensible adhesins contributed to its capacity to interact with the cell lineages tested. Additionally, EC121 remained invasive in the presence of D-mannose but with reduced bacterial counts in T24 cells (P< 0.003) whereas there was no difference in the invasiveness in Caco-2 (Figure 6), showing that mannose-sensible adhesins contributed to both adherence and invasion of EC121, although it was not the only factor associated to these traits.

**Figure 5.**
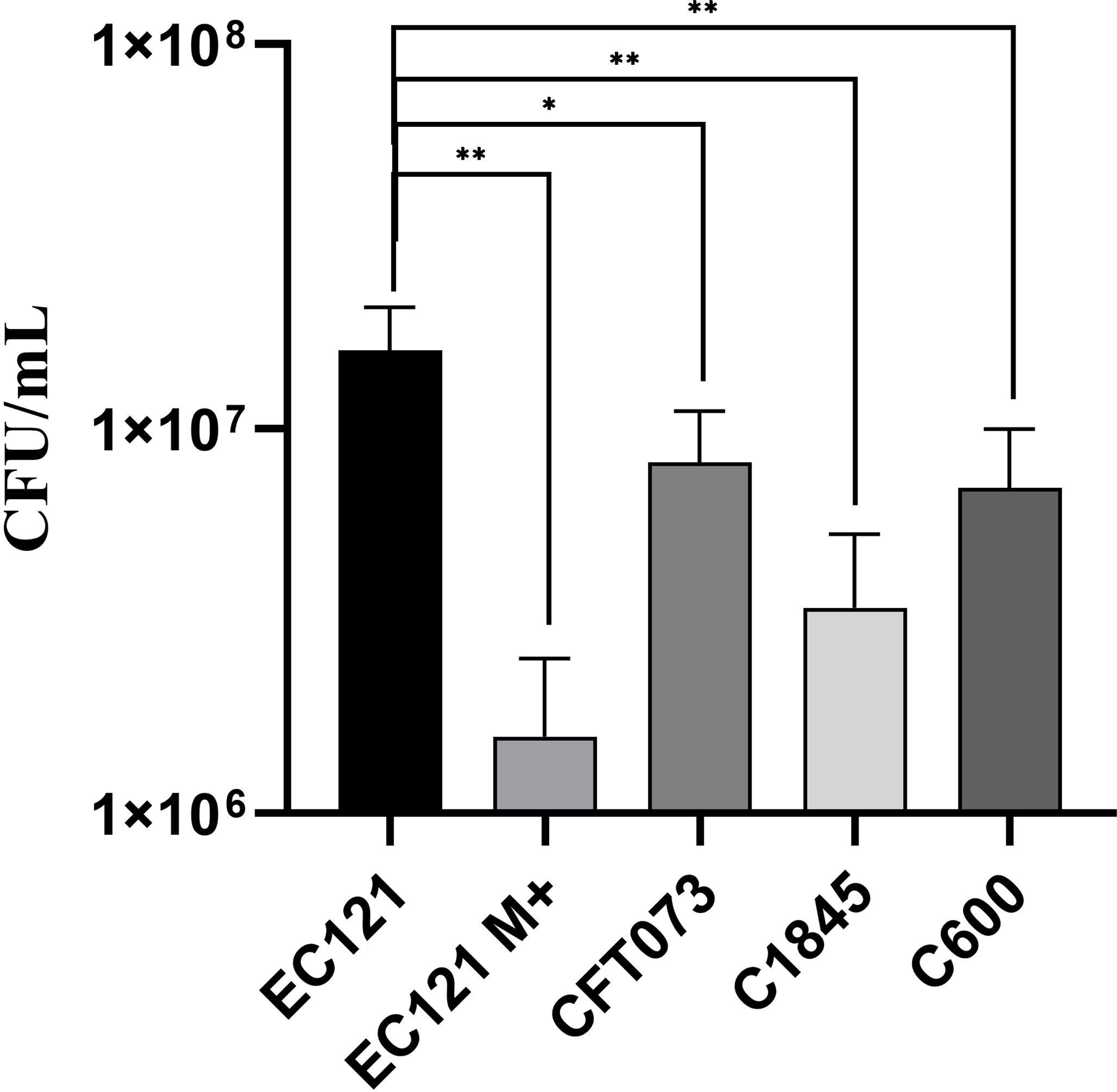
Quantitative adherence assay on HeLa cell. Quantitative assay performed in HeLa cells (3-h of incubation at 37°C) in the absence of D-mannose, except when indicated.; M+, assay performed in the presence of 2% D-Mannose. The CFT073, C1845, and C600 strains were used as controls. Experiments were done in biological duplicates and experimental triplicates. *P* values: * < 0.05, ** < 0.01.

**Figure 6.**
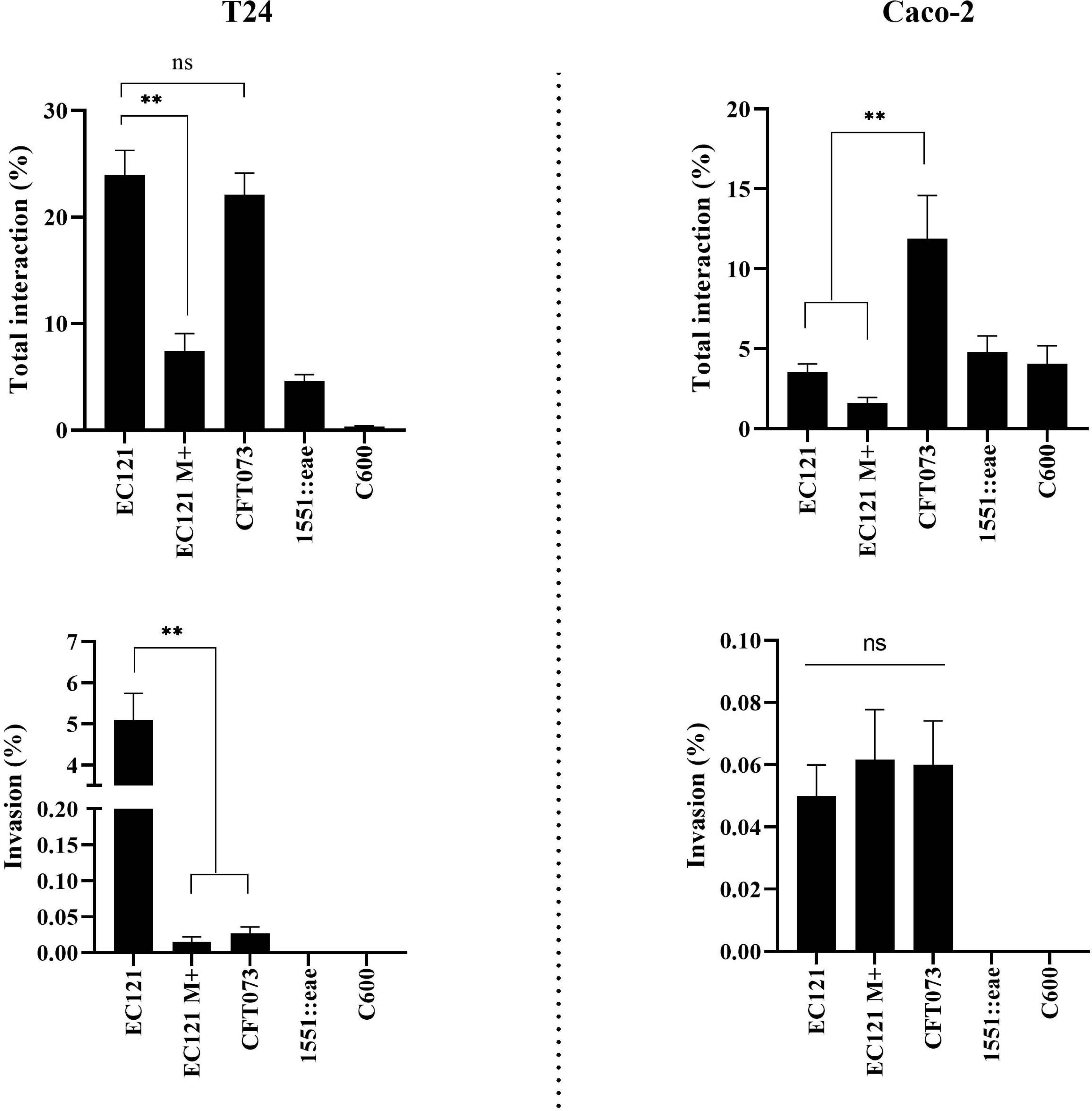
–. Interaction and invasion in different eukaryotic cell lineages**. Short** period quantitative invasion assay was performed in Caco-2 and T24 cells using a MOI of 50 in confluent cell cultures. All assays were carried out in the absence of D-mannose, except for EC121 M+, where 2% D-mannose was added in the media; *E. coli* CFT073 was used as ExPEC control and *E. albertti*1551-2::eae was used as adherent non-invasive control. The assays were performed in experimental triplicates and biological duplicates. *P* values: ** *<*0.008; ns > 0.05

### EC121 is virulent in the *Galleria mellonella* virulence model

The utilization of the *G. mellonella* virulence model showed that EC121 promotes higher mortality rates than the non-pathogenic *E. coli* strain C600 (*P<* 0.005) and the mock-injection (*P*< 0.0001). Although the *E. coli* strain CFT073 killed more larvae than the EC121, there was no significant difference among their survival curves (Figure 7), showing that EC121 is virulent in the model used.

**Figure 7.**
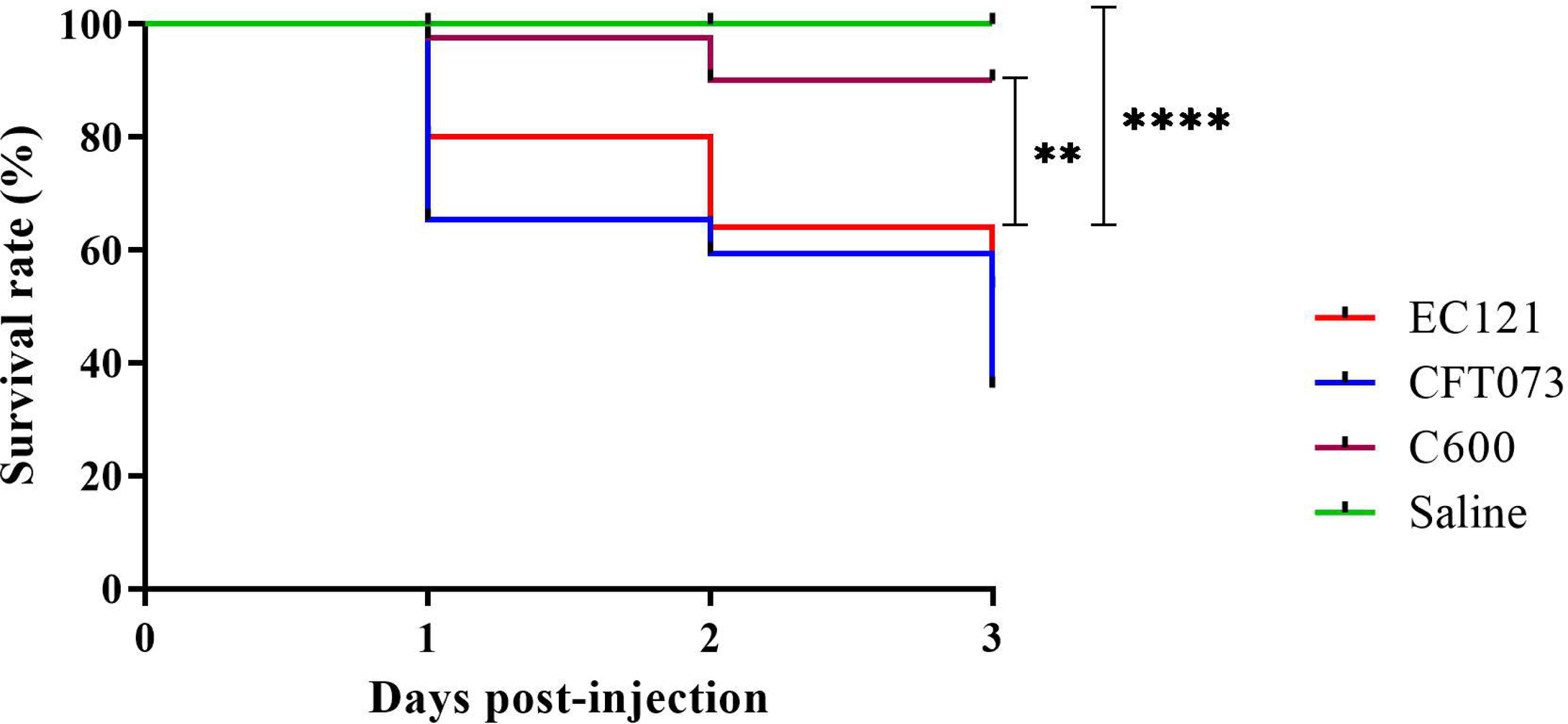
*Galleria mellonella* infection model. The virulence of EC121 was assessed using an inoculum of 10^4^ CFU per *G. mellonella* larvae. The larvae survival rate, expressed by Kaplan-Meier plots indicated that EC121 was more virulent than the negative control (saline) and the non-pathogenic K-12 derived *E. coli* C600, while there was no significant difference between the survival rates of EC121 and the ExPEC prototype strain CFT073. *P* value: ** <0.005; **** <0.0001. The assay was performed three time with five larvae per group.

## Discussion

Authors should discuss the results and how they can be interpreted in perspective of previous studies and of the working hypotheses. The findings and their implications should be discussed in the broadest context possible. Future research directions may also be highlighted. The *E. coli* EC121 strain was isolated in 2007 from a bloodstream infection of an inpatient that presented persistent gastroenteritis and T-zone lymphoma. Since initial analyses showed that it belonged to phylogenetic group B1 and carried few virulence markers commonly related to extraintestinal pathogenic *E. coli,* it was classified as ExPEC negative (ExPEC*-*) [25] and therefore considered as an opportunistic pathogen. However, considering that about 40% of extraintestinal infections are caused by strains devoid of virulence factors [13, 14] and that EC121 was an MDR strain, its entire genome was sequenced to further understand its virulence potential. Interestingly, the EC121 strain belongs to ST101, which has been previously reported to be involved in nosocomial outbreaks caused by Metallo-β-lactamases-producing strains in many countries from Europe, Asia, and Oceania [58–62]. Furthermore, ST101 has also been detected among strains of non-outbreak related extraintestinal infections [63–70], water [71], poultry infection [72], retail food [63,73– 75], and healthy human and animal intestinal microbiota [70,76–79], mostly presenting an MDR phenotype. Shrestha et al. [80] drew attention to ST101 due to the PDR phenotype presented by some strains of this ST, and mainly because it is not considered a pandemic clone, although it has been isolated worldwide. We, therefore, analyzed data about the infection type, isolation source, and resistance genetic markers presented by the strains of the ST101 complex that were previously deposited in the NCBI (Table S1). Such analysis evidenced that MDR strains of this complex were spread worldwide. In addition, such ST complex is related to Shiga toxin-producing strains as well as strains isolated from extraintestinal infections, human and animal microbiota, retail food, and environment. Moreover, many strains simultaneously carry the *bla*_CTX-M-55_, *mcr-1*, *fosA3*, and *qnrS1* genes. Interestingly, one strain present simultaneously *aac(6’)-Ib-cr*, *bla*_CTX-M-55_, *bla*_NDM-5_, *bla*_OXA-1_, *mcr-1*, *fosA3*, and other 19 resistance genes. Likewise, the EC121 strain showed multiple antimicrobial resistance genes, including genes that confer resistance to third-generation cephalosporins (*bla*_CTX-M-2_). It is worth mentioning that the EC121 strain was isolated in 2007, and its MDR phenotype was relevant since at that period it was susceptible only to carbapenems, polymyxins, and amikacin. Recently, the *E. coli* strain ICBEC72H, which belongs to ST101 and carried only *bla*_CTX-M-8_ and *mrc-1*[64] was isolated from a human extraintestinal infection in Brazil. Similarly, the ST101 *E. coli* strain 200H (Table S1) was isolated from a human urinary tract infection and carried *bla*_OXA-9_, *mcr-1*, and *aac(6’)-Ib-cr*. These reports show that MDR strains belonging to the ST101 complex have been circulating in Brazil for a long time. Some authors showed that the *E. coli* strain 912 (ST101) was selected by the usage of antimicrobial agents in animals and that it was able to colonize human and pig gut and spread through the environment, reaching and colonizing animals that were not under antimicrobial treatment [81, 82]. These same authors have also shown that ST101 strains can naturally acquire and transfer plasmid-borne antimicrobial resistance genes in the gut [81, 82]. Such a feature is important for various reasons. First, strain EC121 carried multiple plasmids, which harbored different combinations of antimicrobial-resistance genes, all of which were successfully transferred to *E. coli* K-12 derived strains in a 3-hours conjugation assay. Besides, *E. coli* strains belonging to ST101 were recovered from retail meat in Europe and Asia [74, 75], and from extraintestinal infections in Brazil and the USA in the same regions in which they were detected from retail meat [63, 72]. Additionally, strains belonging to the ST101 complex carrying multiple resistance genes were recovered from the intestine of healthy humans and animals in many countries. Therefore, even if these strains do not cause infection directly, they could potentially transfer plasmids to other bacteria, even from distinct genera. Such cross genera plasmid transfer could be easily identified in the plasmids reported in the present study; IncM1 plasmid, for example, is closely related to plasmids found in *Klebsiella* spp. and *Enterobacter* spp., while IncHI2/HI2A is related to *Salmonella* spp. and *Yersinia* spp. plasmids. Together, these findings reinforce the high risks associated with strains belonging to the ST101 complex due to their ability to colonize humans and animals’ gut, to easily disseminate via retail food and water, being able to acquire and spread antimicrobial resistance-encoding genes. Strains from the ST101 complex are included in the phylogenetic group B1, which implies that they do not have all the classical virulence factors that are usually associated with the most virulent ExPEC strains [83, 84]. Many studies reported phylogroup B1 *E. coli* strains as commensals or as intestinal pathogens, but not as extraintestinal pathogens [83–85]. The genomic analysis of the EC121 strain showed a high number of virulence genes, demonstrating that it presents all the traits necessary to be considered as an extraintestinal pathogenic agent, like adhesins, iron acquisition systems, and genes related to immune system evasion. Moreover, like other ExPEC strains, EC121 displayed multiple virulence genes related to each feature, reflecting the redundant phenotype that ensures its pathogenicity. However, even considering the completeness of each sequence and each operon, which was manually checked, the presence of virulence genes “per se” does not guarantee that all of them are expressed. Therefore, to evaluate the expression of such traits, distinct phenotypic assays were performed and confirmed the virulence genetic background of EC121. To test the bacterial ability to resist the serum complement activity that could be conferred by the presence of *traT, iss,* and *ompT*, a two hours challenge assay was performed, in which one particle can traverse all circulatory system at least twice, so a pathogen that resists complement’s activity during this period, even with a small bacterial load, is, in theory, more capable of reaching different niches and spread through the bloodstream or cause a bloodstream infection. The EC121 strain resisted the NHS for two hours with an inoculum similar to the resistant *E. coli* J96 control strain, thus confirming the EC121 serum resistant phenotype. Serum complement is the first immunological barrier to control pathogens that reach the bloodstream. Complement resistance confers the possibility to spread to different body sites through the bloodstream. Hallström et al. [86] reported the relationship of bacterial resistance to NHS with sepsis severity, and other authors have associated it with different kinds of extraintestinal infections [87–89]. The ability to colonize and attach to surfaces is also an important trait for any pathogenic bacteria; in this way, the assays carried out showed not only that EC121 strain was able to adhere to and invade different cell lineages, including bladder T24 cells but that it could also attach and produce biofilm on abiotic surfaces. Peirano et al. [90] showed that ExPEC negative ST101 MDR strains isolated from extraintestinal infections could interact with HEp-2 and Caco-2 cells more efficiently than strains belonging to the epidemic clones ST131 and ST405, which are ExPEC positive [90]. The strain EC121 interacted with all cell lineages tested, but its interaction was significantly higher in T24 cells. Besides, its interaction’s capacity was similar to the strain CFT073 (ExPEC prototype), and it invaded T24 cells more efficiently, suggesting that EC121 might be capable to produce intracellular bacterial communities (IBC). IBCs were related to bacterial persistence and recurrent infections in the host [91]. Overall, the EC121 invasiveness might be more related to its persistence ability than to its capacity to transpose epithelial barriers, since persistence may help bacteria to evade the immune system and confer protection against antimicrobials agents. However, further studies are required to prove this hypothesis. Ten virulence encoding-genes involved with biofilm production were identified in the EC121 genome (Table S3). The capacity to produce biofilm might confer many advantages to bacteria, like protection against the immunological system and antibiotics, assisting its persistence and spread in the environment. However, biofilm production depends on many factors, like temperature and presence of specific nutrients in the media or environment [92–94]. Moreover, the ability to produce biofilm has been reported to vary among ExPEC strains and that strains with this capacity are considered to be more pathogenic [95].

Interestingly, many of the EC121 virulence factors detected in the draft genome are related to diarrheagenic *E. coli*, even though none of them is implicated in DEC pathotype definition. The presence of many genetic features related to Shiga-toxin producing *E. coli* (STEC) strains, *e.g*., Hcp, EhaG, and Lpf-1_O26_, as well as the proximity of EC121 to the clade that contains STEC strains and *E. coli* O104:H4 strain 2011C-3493, draws attention to its potential diarrheagenic background. Many features identified in the EC121 genome reinforce its linkage with STEC strains. Phenotypically, EC121 expressed the O154:H25 serotype, but it possesses the group IV capsule-encoding genes. This kind of capsular group is known to be thermoresistant and expressed as K_LPS_ or O-antigen capsule. This could explain the expression of the O154 instead of O100 antigen, despite the presence of all genes related to the expression of the latter. Interestingly O100 is a STEC related serogroup. Moreover, much of the phage remains detected in the EC121 strain were related to Stx-converting phages; besides, ST101 strains carrying the *stx_1a_* gene have been reported in food sources [73, 96]. In humans, ST101 strains were already reported in a patient with Hemolytic Uremic Syndrome (HUSEC) [96–98] and in non-bloody diarrhea related to a Stx1a-producing *E. coli* strain [99]. Although only one of these strains had its genome sequenced, some ST101 *E. coli* strains recovered from animals and food were found to carry *stx1.* Interestingly, most Stx-converting phages remains found in EC121 were similar to those commonly related to Stx1a production, corroborating with the results presented here. The genome of three non-STEC strains from diarrheic patients was found in GenBank, one of which was devoid of DEC virulence factors. Likewise, EC121 was isolated from bloodstream infection of one inpatient with persistent infectious gastroenteritis which was probably the source of EC121 infection. Unfortunately, the *E. coli* isolated from stool was not stored not allowing further comparison between bloodstream and stool *E. coli* isolates. Several studies evaluated *E. coli* pathogenicity in the surrogate *G. mellonella* model [56,100–104], these studies point to the model efficiency in differentiating pathogenic strains from non-pathogenic, especially using bacterium inoculum of 10^5^ CFU per larvae or lower [103, 104]. Moreover, Jonsson et al. [100] have shown that pathogenic *E. coli* needs an inoculum of at least 10^3^ CFU to lead larvae to death. To evaluated EC121 virulence, a bacterial inoculum of 10^4^ CFU per larvae was used, and in the assayed conditions EC121 was as virulent as the ExPEC prototype CFT073, thus corroborating with the *in silico* and *in vitro* data, showing that some strains from the ST101 are truly pathogenic.

In summary, our extensive *in silico, in vitro,* and *in vivo* analyses of virulence and resistance properties of *E. coli* strain EC121, an O154:H25 B1-ST101 strain isolated from a human bloodstream infection, confirmed its virulence potential and increased the knowledge on the complex scenario of virulence traits present in the MDR *E. coli* ExPEC negative group, contributing to the potential development of strategies to control the spread of such pathogens.

## Acknowledgments

We are grateful to the Laboratório Especial de Microbiologia Clínica (LEMC) of the Federal University of São Paulo (UNIFESP), São Paulo - SP, Brazil, for providing the *E. coli* strain EC121.

## Ethics

The strain EC121 used in this research was obtained from clinical routine after laboratory procedures. No additional procedure was performed to acquire any bacterial strain, so the consent form was not required as determinate by the Brazilian National Health Council n° 466/12 and 510/16. All patient information was obtained from medical records, and the research was done with the approval of the local Research Ethics Committee of the Federal University of São Paulo - UNIFESP/São Paulo Hospital (CEP 2031/08 and CEP N 7140160317).

## Supporting information captions

**Table S1** - ST101 complex strains, NCBI accession, multidrug resistance genotype, source, and country of isolation.

**Table S2** - Type V Secretion system protein identified by MaSyFinder.

**Table S3** - Complete Genetic cluster identified in EC 121 strain and their predicted association with virulence.

**Table S4** - Virulence genes identification and validation in NCBI and Swiss-Prot.

**Table S5** - Detailed information of all predicted phages detected in strain EC121.

**Figure S1** - Phylogenetic relationship among 62 human isolates from the ST101 complex

**Figure S2** - Plasmid content of the EC121strain.

**Figure S3** - Phylogenetic relationship obtained from sequence alignment of pEC121_IncHI2/HI2A, using BLASTn

**Figure S4** - Phylogenetic relationship obtained from sequence alignment of pEC121_IncM1 plasmid, using BLASTn

**Figure S5** - EC121 adherence in eukaryotic cell linages HeLa, Caco-2, and T24.

